# Stomatal, mesophyll, and biochemical limitation to photosynthesis of soybeans under waterlogging and reoxygenation

**DOI:** 10.1101/2025.10.17.683018

**Authors:** Shigehiro Kubota, Gaku Yokoyama, Toshimune Shibata, Daisuke Yasutake, Tomoyoshi Hirota

## Abstract

While waterlogging stress slows photosynthetic rate (*A*_sat_), the underlying processes remain poorly understood. Here, we aimed to characterize the limitations to photosynthesis imposed by stomatal conductance (*g*_s_), mesophyll conductance (*g*_m_), and biochemical processes under waterlogging and subsequent reoxygenation. Two soybean cultivars (*Glycine max* L. cv. Fukuyutaka and Iyodaizu) were subjected to 6 days of waterlogging, after which excess water was drained. The responses of *A*_sat_, *g*_s_, *g*_m_, and the maximum carboxylation rate (*V*_cmax_) were investigated. In both cultivars, *A* declined significantly within 4 days of waterlogging and did not recover completely by two weeks of reoxygenation. During waterlogging, CO_2_ concentration at carboxylation site decreased in parallel with *g*_s_ and *g*_m_, indicating that photosynthesis was mainly limited by diffusional factors (combination of *g*_s_ and *g*_m_). After drainage, diffusional limitation persisted during early reoxygenation, whereas biochemical limitation due to reduced *V*_cmax_ became dominant after 7 days of reoxygenation. Therefore, maintaining high diffusional conductances and *V*_cmax_ during waterlogging and reoxygenation, respectively, is important for enhancing photosynthetic tolerance to waterlogging stress. Overall, our results demonstrate that *A*_sat_ under waterlogging and reoxygenation is dynamically constrained by multiple factors, emphasizing the need for comprehensive assessment of gas diffusion and carbon assimilation processes.

## 1 INTRODUCTION

More than 17 million km^2^ of land area is exposed to waterlogging, caused by precipitation or irrigation exceeding soil drainage capacity, worldwide every year (Voesenek and Sasidharan, 2013). The spatiotemporal precipitation pattern is expected to change as climate change progresses, thereby increasing the risk of waterlogging in some regions (Hirabayashi et al., 2013). Because oxygen diffusivity in water is 10^4^-fold smaller than that in air, soil waterlogging causes oxygen depletion in the rhizosphere, resulting in restricted plant growth and yield (Herzog et al., 2016; Manghwar et al., 2024). This inhibition of plant growth imposed by waterlogging is mainly attributed to the reduction of photosynthetic CO_2_ assimilation (Manghwar et al., 2024; Ploschuk et al., 2018). The photosynthetic process comprises gas diffusion from ambient air to chloroplast and carbon assimilation reactions which are controlled by stomatal and mesophyll conductance and the maximum carboxylation rate and maximum electron transport rate, respectively (Grassi and Magnani, 2005). Although photosynthetic responses to waterlogging have been widely investigated for decades to understand plant adaptation and survival strategies (Herzog et al., 2016; Mommer and Visser, 2005), the primary limiting factor of photosynthesis remains unclear.

Waterlogging impedes aerobic respiration in the roots via oxygen depletion, inducing the depletion of root ATP levels, acidification of cytosol, and an increase in reactive oxygen species (ROS, ·O_2_ ^−^ and H_2_ O_2_) (Garcia et al., 2020; da-Silva and Amarante, 2020). These chemical signals inhibited the expression and activity of aquaporins (Aroca et al. 2012; Tournaire-Roux et al., 2003), leading to a reduction in root hydraulic conductance and a subsequent decrease in leaf water potential (Else et al., 2001; Garcia et al., 2020; Kubota et al., 2025; Toral-Juarez et al., 2021). As a result, despite sufficient available water in the soil, waterlogging triggers leaf water stress (Haverroth et al., 2025; Kubota et al., 2025). Given that stomatal closure acts as the initial and most prominent constraint on photosynthesis under soil drought (Flexas et al., 2009; Grassi and Magnani, 2005; Galle et al., 2009), stomatal conductance could also represent a primary limiting factor under waterlogging. Indeed, the photosynthetic rate decreases in coordination with stomatal conductance during the early period of soil waterlogging in some plant species (Ploschuk et al., 2018; Toral-Juarez et al., 2021).

In contrast, some recent studies have pointed out that a decrease in the photosynthetic rate cannot be exclusively explained by a decrease in stomatal conductance (Herzog et al., 2016, Olorunwa et al., 2022b, Ploschuk et al., 2018). For instance, in barley, the decrease in photosynthetic rate under waterlogging was attributed to a reduction in mesophyll conductance (Ploschuk et al.,

2018). In addition, prolonged waterlogging causes the accumulation of leaf ROS (da-Silva and Amarante, 2020), resulting in oxidative stress. This stress induces the impairment of ribulose-1,5-bisphosphate carboxylase/oxygenase (Rubisco) activity (Olorunwa et al., 2022a), the loss of photosynthetic pigments (Garcia et al., 2020; da-Silva and Amarante, 2020), and damage to the photosynthetic electron transport chain (Posso et al., 2020; Martins et al., 2024). As a result, photosynthetic capacity (maximum carboxylation rate and maximum electron transport rate) decreases under soil waterlogging for more than one week (Lee et al., 2022; Yang et al., 2024), indicating the biochemical photosynthetic limitation under prolonged waterlogging. Thus, the limiting factor of photosynthesis under waterlogging may change over time and involve not a single physiological process but rather multiple factors acting simultaneously.

Interestingly, leaf gas exchange rate rarely recovers immediately after the drainage of excess water; instead, plants are exposed to further stress conditions (Garcia et al., 2020; Toral-Juarez et al., 2021). A drastic increase in oxygen supply to the root by reoxygenation causes redox imbalance and overproduction of ROS in both roots and leaves (da-Silva and Amarante, 2020), which induces oxidative stress. Thus, soil reoxygenation can enhance the decrease in maximum photosynthetic capacity (Toral-Juarez et al., 2021), indicating a biochemical limitation to photosynthesis. Moreover, leaf water depletion due to hydraulic dysfunction continues after waterlogging (Haverroth et al., 2025; Kubota et al., 2025), which induces a decrease in stomatal and mesophyll conductance. As the recovery of photosynthesis from waterlogging-induced stress is essential for resuming plant growth, examining the photosynthetic response during post-waterlogging reoxygenation is important for understanding the acclimation mechanism and evaluating plant tolerance (Martins et al., 2024).

A framework to analyze photosynthetic limitation processes was introduced by Grassi and Magnani (2005), who partitioned the total constraint on net photosynthesis into three distinct components: stomatal conductance, mesophyll conductance, and biochemical processes (i.e. carboxylation activity). Since then, quantitative limitation analysis has been widely applied to evaluate changes in the relative proportions of photosynthetic limitations under soil drought (Galle et al., 2009; Flexas et al., 2009; Egea et al., 2011). However, this has not been applied to waterlogging-stressed plants so far. As described, waterlogging causes leaf dehydration via a decrease in plant hydraulic conductance, and thus photosynthetic limitation dynamics under waterlogging may exhibit similar patterns to those observed under drought. In contrast, given that the direct cause of waterlogging stress is inhibition of root respiration, different mechanisms (e.g., oxidative stress due to accumulation of ROS and active stomatal closure due to accumulation of ABA) may affect the photosynthetic limitation dynamics (Haverroth et al., 2025). Because the comprehensive measurement of temporal changes in stomatal and mesophyll conductance and photosynthetic capacity during waterlogging and reoxygenation are scarce (Yang et al., 2024), the primary limitation factors attributing such declines in photosynthetic rate remain poorly understood.

The objective of this study was to evaluate the stomatal, mesophyll, and biochemical limitations to photosynthesis during waterlogging and subsequent reoxygenation. We hypothesized that the photosynthetic rate is primarily constrained by stomatal limitations during the early period of waterlogging, whereas the proportions of mesophyll and biochemical limitations increase from late waterlogging through reoxygenation. To test the hypothesis, we measured gas exchange and chlorophyll fluorescence responses of two soybean cultivars (*Glycine max* (L.) Merr., cv. Fukuyutaka and Iyodaizu) to soil waterlogging and reoxygenation and conducted quantitative limitation analysis. These cultivars have contrasting growth strategies under waterlogging conditions; Fukuyutaka suppresses the growth during waterlogging, while Iyodaizu maintains a high growth rate (Sakazono et al., 2013; Suematsu et al., 2017), which is expected to show contrasting response of photosynthetic rate to waterlogging stress.

## 2 MATERIALS AND METHODS

### 2.1 Plant materials, growth conditions, and experimental design

The study was conducted in a greenhouse located at the Ito Plant Experiment Fields & Facilities, Faculty of Agriculture, Kyushu University, Fukuoka, Japan (33°35.5’N, 130°12.9’E). A ventilation fan was operated when the air temperature exceeded 20 °C, and the side-wall windows of the greenhouse were kept open during the cultivation period. We chose two soybean cultivars: Fukuyutaka and Iyodaizu (Fig. S1 (A)). Fukuyutaka is the most popular cultivar in Japan, and Iyodaizu is a waterlogging-tolerant cultivar, which can maintain a high growth rate under waterlogging (Sakazono et al., 2013; Suematsu et al., 2017). Seeds (*n* = 60) of each cultivar were sown in 0.11-L polyethylene pots filled with a soil mixture of vermiculite and decomposed granite in a volume ratio of 5:4 on 9 May 2023. The seedlings were irrigated as needed with tap water until the unifoliolate leaves fully expanded (VC stage). Seedlings (*n* = 20) of each cultivar were then transplanted into 8 L pots (23 cm height and 30 cm diameter) filled with the same soil used for cultivating seedlings on 22 May. Plants were irrigated with a nutrient solution (Otsuka AgriTechno Co. Ltd., Japan) with an electrical conductivity of 0.8 dS m^−1^ one to three times per day. The nutrient solution contained 17.1 mmol (NO_3_ ^−^) L^−1^, 1.1 mmol (PO_4_ ^3−^) L^−1^, 1.6 mmol (SO_4_ ^2−^) L^−1^, 8.4 mmol (K^+^) L^−1^, 1.5 mmol (Mg^2+^) L^−1^, and 3.9 mmol (Ca^2+^) L^−1^. As soybean grew, the concentration of solution was increased to 1.2 dS m^−1^ on 12 June. Air temperature (*T*_a_), and relative humidity (RH) in the greenhouse were measured using a temperature/humidity sensor with a built-in data logger (MX2302A, Onset Computer Corp., Massachusetts, USA). Solar radiation (*I*) outside the greenhouse was obtained from a meteorological station (pyranometer: CHF-SE20-JM, Climatec Inc., Tokyo, Japan) located about 170 m away from the greenhouse. The data were measured at 10-s intervals, and the mean values were logged at 10-min intervals. Measured data are shown in Fig, S2.

Soil waterlogging was initiated on 14 July when soybeans were anthesis stage (R1 stage). Soybean plants of each cultivar were randomly divided into two groups (*n* = 5): control and waterlogging. In waterlogging treatment, two cultivation pots were placed into each of five large containers (60 L; 22.5 cm height, 63 cm length, and 42.5 cm width), and subsequently all containers were watered from the bottom of the container at 7:00, taking care not to trap air bubbles in the bottom of pots (Fig. S1 (B)). We watered every one to three days to maintain the water level at least 2 cm above the soil surface. To investigate the recovery process after drainage, the excess water was rapidly drained at 7:00 on 20 July. In control treatment, plants were irrigated three times per day.

### 2.2 Measurements of leaf gas exchange and chlorophyll fluorescence

Leaf gas exchange and chlorophyll fluorescence measurements were performed with two portable photosynthesis systems (Li-6400XT, LI-COR, Lincoln, NE, USA) equipped with a leaf chamber fluorometer (LI-6400-40, LI-COR, Lincoln, NE, USA). Light-saturated photosynthetic rate (*A*_sat_), stomatal conductance for water vapor (*g*_sw_), intercellular CO_2_ concentration (*C*_i_) and chlorophyll fluorescence were measured from 7:00 to 10:00. The fully expanded leaves exposed to direct sunlight were selected for measurements. The leaves were clamped at least 15 min and allowed to achieve steady-state gas exchange. Leaf gas exchange was measured with saturating light (1400 μmol m^−2^ s^−1^). Air temperature inside the chamber was controlled to follow ambient conditions from 28 to 33 °C, and vapor pressure deficit was controlled so as not to exceed 2.0 kPa as possible. As the one-point method can provide inaccurate estimates of maximum carboxylation rate (*V*_cmax_) when photosynthesis is not RuBP-saturated, Burnett et al. (2019) recommended a measurement at *C*_a_ values below the current ambient CO_2_ concentration. Thus, the CO_2_ concentration inside the chamber (*C*_a_) was set to 350 μmol mol^−1^ for the stable estimation of *V*_cmax_.

From the fluorescence measurements, the photochemical efficiency of photosystem □ (Φ_PS□_) was determined as follows:

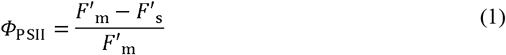

where *F*_m_ is the maximum fluorescence during a light-saturated pulse (∼7200 μmol m^−2^ s^−1^), and *F*’_s_ is the steady-state fluorescence yield at a PPFD of 1400 μmol m^−2^ s^−1^. Since Φ_PS□_ indicates the ratio of electrons transferred to the absorbed photons by photosystem □, the electron transport rate (*ETR*) was calculated as follows:

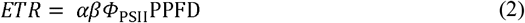

where *α* is the leaf absorptance and *β* is the partitioning fraction of absorbed quanta between two photosystems. Following previous studies, *α* and *β* were assumed to be 0.85 (Evans & Loreto, 2000) and 0.5 (Laisk & Loreto, 1996), respectively. Furthermore, to evaluate the severity of oxidative stress, we calculated *ETR*/*A*_sat_. The increase in *ETR*/*A*_sat_ implies that the light energy exceeds the electron requirements for the photosynthetic carbon assimilation, suggesting that an excess of reducing power that may trigger photoinhibition and photooxidative effects (Baker, 2008; Toral-Juarez et al., 2021).

To derive reliable values of *V*_cmax_ and mesophyll conductance (*g*_m_), the accurate measurements of *A* and *C*_i_ is particularly important (Pons et al., 2009). When *g*_sw_ shows lower values due to soil waterlogging, the contribution of cuticular transpiration to the observed transpiration relatively increases, resulting in an overestimation of *g*_sw_ and *C*_i_ (Márquez et al. 2021). Thus, we measured cuticular conductance (*g*_cut_) after the experiment according to Sack and Scoffoni (2010). Briefly, three leaves for both cultivars were sampled by cutting at the base of petiole from each treatment, and bench-dried for at least 1 hour. Then, the leaves were weighed at intervals of 1−5 min with electronic-balance (resolution 0.001 g; TW423N, Shimadzu Corp., Kyoto, Japan) 15 to 24 times. Air temperature and relative humidity were measured by a temperature–humidity probe (HMP-60, Vaisala, Helsinki, Finland) during the weight measurement. The leaf images were acquired by flatbed scanner (GT-S650, Seiko Epson Corp., Nagano, Japan) before and after the weight measurement, and leaf area was estimated using ImageJ *v*. 1.54 software. We calculated *g*_cut_ using Microsoft Excel (Microsoft Corp., Redmond, Washington, USA) spreadsheet provided by Sack and Scoffoni (2010). As there was no significant difference in *g*_cut_ between treatments, the average value for each cultivar was used for analysis (Table S2). Moreover, the coefficients of diffusional leakage through the gaskets were determined following Flexas et al. (2007). Based on the *g*_cut_ and diffusional leakage, we recalculated *A*_sat_, *g*_sw_, and *C*_i_, and used the corrected values in all the following analyses.

### 2.3 Calculation of mesophyll conductance and biochemical capacity parameters

We estimated *g*_m_ by the variable *J* method (Harley et al., 1992) and *V*_cmax_ by the one-point method (De Kauwe et al., 2016). From the measured data of gas exchange and chlorophyll fluorescence, *g*_m_ was calculated as follows:

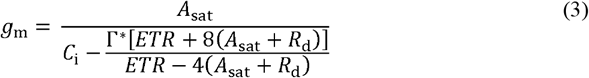

where *R*_d_ is mitochondrial respiration rate in the light, and *Γ*^*^ is the CO_2_ compensation point in the absence of mitochondrial respiration. *R*_d_ was assumed to be 1.5% of *V*_cmax_ at the intercellular CO_2_ concentration (De Kauwe et al., 2016; Oliveira et al., 2023). *A*_sat_ and *C*_i_ were derived from gas exchange measurements, and *ETR* was calculated using Eq. 2. The CO_2_ concentration inside the chloroplast (*C*_c_) was calculated as follows:

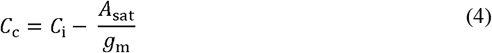

Based on the one-point method proposed by Wilson et al. (2000) and verified by De Kauwe et al. (2016) and Burnett et al. (2019), *V*_cmax_ was estimated from the measured data of gas exchange under light-saturated conditions as follows:

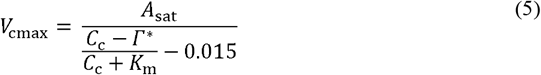

where *K*_m_ is given by

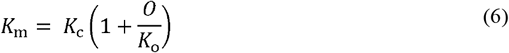

where *K*_c_ and *K*_o_ are the Michaelis-Menten constants for carboxylation and oxygenation, respectively, and *O* is the O concentration (210 mmol mol^−1^). *K*_c_, *K*_o_, and *Γ*^*^ were estimated at the measuring *T*_leaf_ by Arrhenius function and kinetic constants proposed by Bernacchi et al. (2001). Moreover, we scaled *g*_m_ and *V*_cmax_ to 25 °C using a modified Arrhenius function as follows:

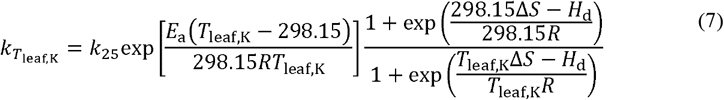

where *k*_Tleaf,K_is the parameter value at a given leaf temperature, *k*_25_ is the parameter value at 25 °C, *E*_a_ is the activation energy, Δ*S* is the entropy factor; *H*_d_ is the deactivation energy, *T*_leaf,K_ is leaf temperature in Kelvin, *R* is the universal gas constant. These parameters (*k*_25_, *E*_a_, Δ*S*) were determined by fitting Equation (7) to the relationship between *T*_laef_ and *V*_cmax_ obtained from the full *A*-*C*_i_ curve method before the waterlogging treatment (Table S1 and Fig. S3). For these curves, the sequence of reference CO_2_ concentration set-points was 400, 300, 200, 150, 100, 50, 400, 600, 800, 1000, 1500 μmol mol^−1^, and four curves were constructed for each leaf with increasing *T*_laef_ from 23 °C to 35 °C with saturated light conditions (1400 μmol m^−2^ s^−1^) and relatively moist air (VPD < 2 kPa as possible). *V*_cmax_ at each leaf temperature was determined using *A*-*C*_i_ curve-fitting utility developed by Moualeu-Ngangue et al. (2017). In addition, we tested estimation accuracy of one-point method by comparing the estimated *V*_cmax_ values with those obtained from the full *A*-*C*_i_ curve method (Fig. S4 and Table S3).

### 2.4 Quantitative limitation analysis of photosynthesis

To evaluate the limitations imposed by waterlogging and reoxygenation on photosynthesis, a quantitative limitation analysis of photosynthesis was performed for all measured data sets according to Grassi and Magnani (2005). According to their approach, the relative changes in light-saturated photosynthetic rates were separated into the three major components: stomatal conductance, mesophyll conductance, and biochemical capacity:

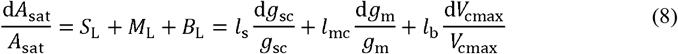

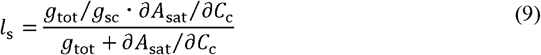

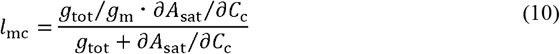

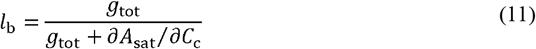

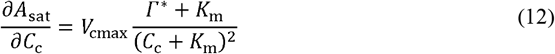

where *g*_sc_ is stomatal conductance to CO_2_ (*g*_sw_/1.6), *g*_tot_ is total conductance to CO_2_ between the leaf surface and carboxylation sites ([1/*g*_sc_ + 1/*g*_m_]^−1^), *S*_L_, *MC*_L_ and *B*_L_ are the contributions of stomatal conductance, mesophyll conductance, and maximum carboxylation rate at 25°C to relative change in photosynthetic rate, respectively, and *l*_s_, *l*_mc_, *l*_b_ are the corresponding relative limitations, with values ranging from zero to one. Then, d*A*_sat_/*A*_sat_ was defined as follows:

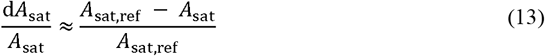

where *A*_sat,ref_ was reference value of *A*_sat_. In the current study, *A*_sat_ of control treatment was used as reference. d*g*_sc_/*g*_sc_, d*g*_m_/*g*_m_, and d*V*_cmax_/*V*_cmax_ were determined in a similar way in equation (13).

### 2.5 Statistical analysis

We utilized five plants of each cultivar in each of the two watering conditions (*n* = 5). As one-point method accurately estimates *V*_cmax_ only when photosynthesis is RuBP-saturated (Burnett et al., 2019), we excluded the data in which *C*_c_ was below the transition point from Rubisco limitation to RuBP limitation in photosynthetic assimilation. The transition point of *C*_c_ (*C*_ctr_) was calculated as follows:

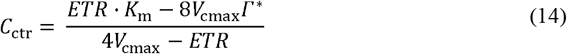

Similarly, we excluded the data in the range of 10 < d*C*_c_/d*A*_sat_ < 50 for accurate analysis of *g*_m_ according to previous studies (Harley et al., 1992), where d*C*_c_/d*A*_sat_ was calculated as follows:

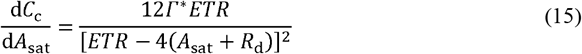

Thus, the replicate number was three to four in some cases. *Student*’s *t*-tests were performed to test differences between control and waterlogged plants of each cultivar. To analyze the relationship between *A*_sat_ and photosynthetic parameters (*g*_s_, *g*_m_, *V*_cmax_, *ETR, C*_i_, *C*_c_), *Spearman*’s rank correlation coefficient was calculated, and the significance of the correlation was confirmed. These analyses were performed using R software (version 4.4.2, R Development Core Team, Vienna, Austria).

## 3 RESULTS

### 3.1 The response of leaf gas exchange to soil waterlogging and reoxygenation

In both cultivars, *A*_sat_ and *g*_sc_ under waterlogging significantly decreased compared to the control after 4 days of waterlogging (Fig. 1). In Fukuyutaka, *A*_sat_ and *g*_sc_ decreased to 40% and 16% of the control on 6 days of waterlogging and recovered to 77% and 97% of the control 7 days after reoxygenation (13 days of total treatment), respectively (Fig. 1 (A) and (C)). In Iyodaizu, *A*_sat_ and *g*_sc_ decreased to 33% and 17% of the control on 6 days of treatment and partially recovered to 54% and 65% of the control at the end of the experiment, respectively (Fig. 1 (B) and (D)). The *g*_m_ and *C*_c_ also showed significant reductions during late-waterlogging period and recovered coordinately during post-waterlogging reoxygenation in both cultivars (Fig. 1 (E) to (H)). In contrast, *V*_cmax_ in the waterlogging was significantly decreased compared to the control only during the reoxygenation period (Fig. 2 (A) and (B)). *ETR* was significantly decreased from the 4 days of treatment and hardly recovered during reoxygenation (Fig. 2 (C) and (D)). *ETR*/*A*_sat_ sharply increased immediately after drainage and remained higher than the control throughout the reoxygenation period in both cultivars (Fig. 2 (E) and (F)).

**Fig. 1:**
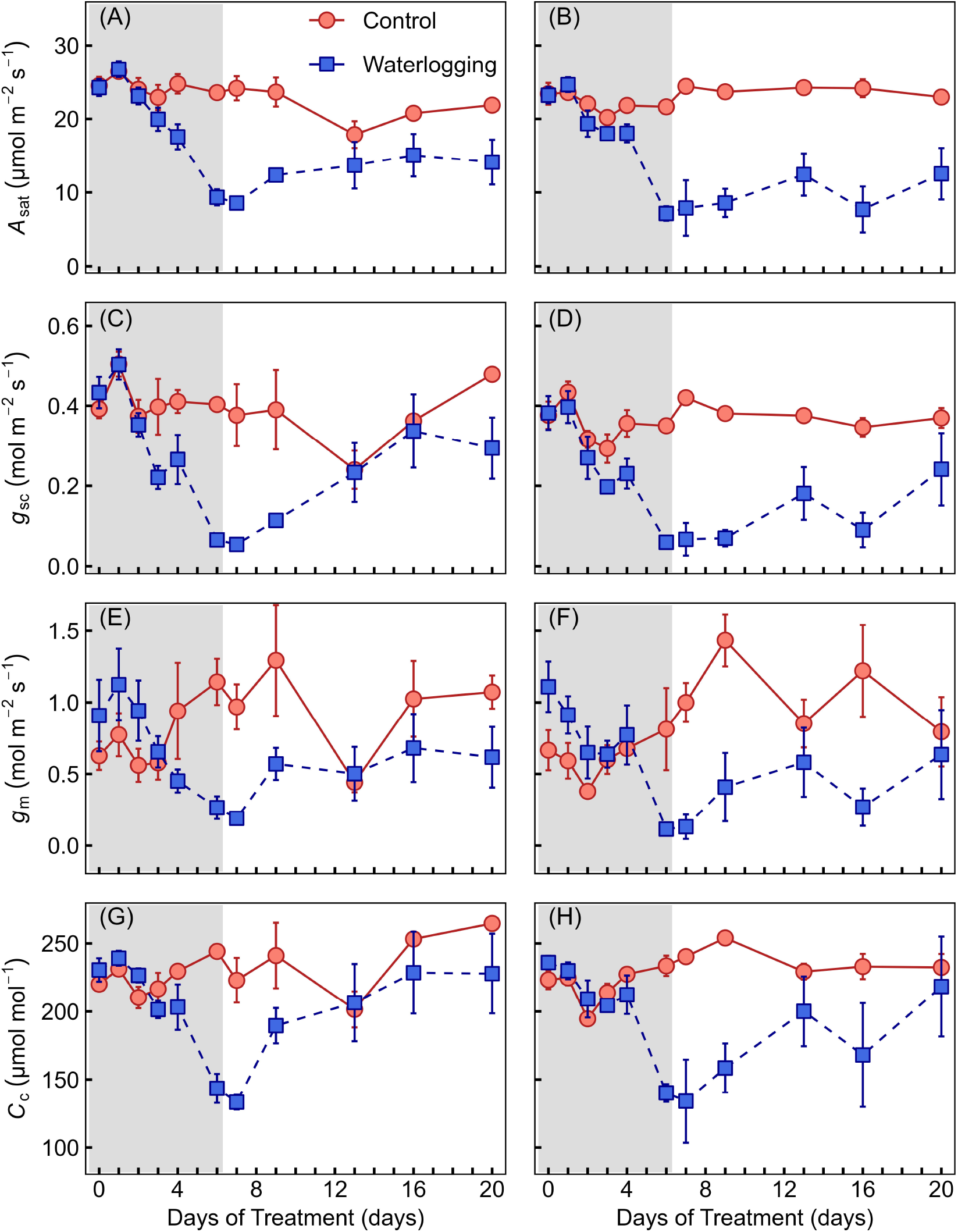
Temporal changes in light-saturated photosynthetic rate *A*_sat_ (A, B), stomatal conductance for CO_2_ *g*_sc_ (C, D), mesophyll conductance *g*_m_ (E, F), and CO_2_ concentration at the carboxylation site *C*_c_ (G, H) during waterlogging and reoxygenation period. The red circles and solid line indicate the measured values in control, and the blue squares and dushed line indicate the measured values in waterlogging treatment. The respective rows indicate the soybean cultivar (i.e. A, C, E, G: Fukuyutaka, B, D, F, H: Iyo-daizu). Gray shaded area indicates the waterlogging period. Plots and error bars denoted means ± standard error (*n* = 4 to 5).

**Fig. 2:**
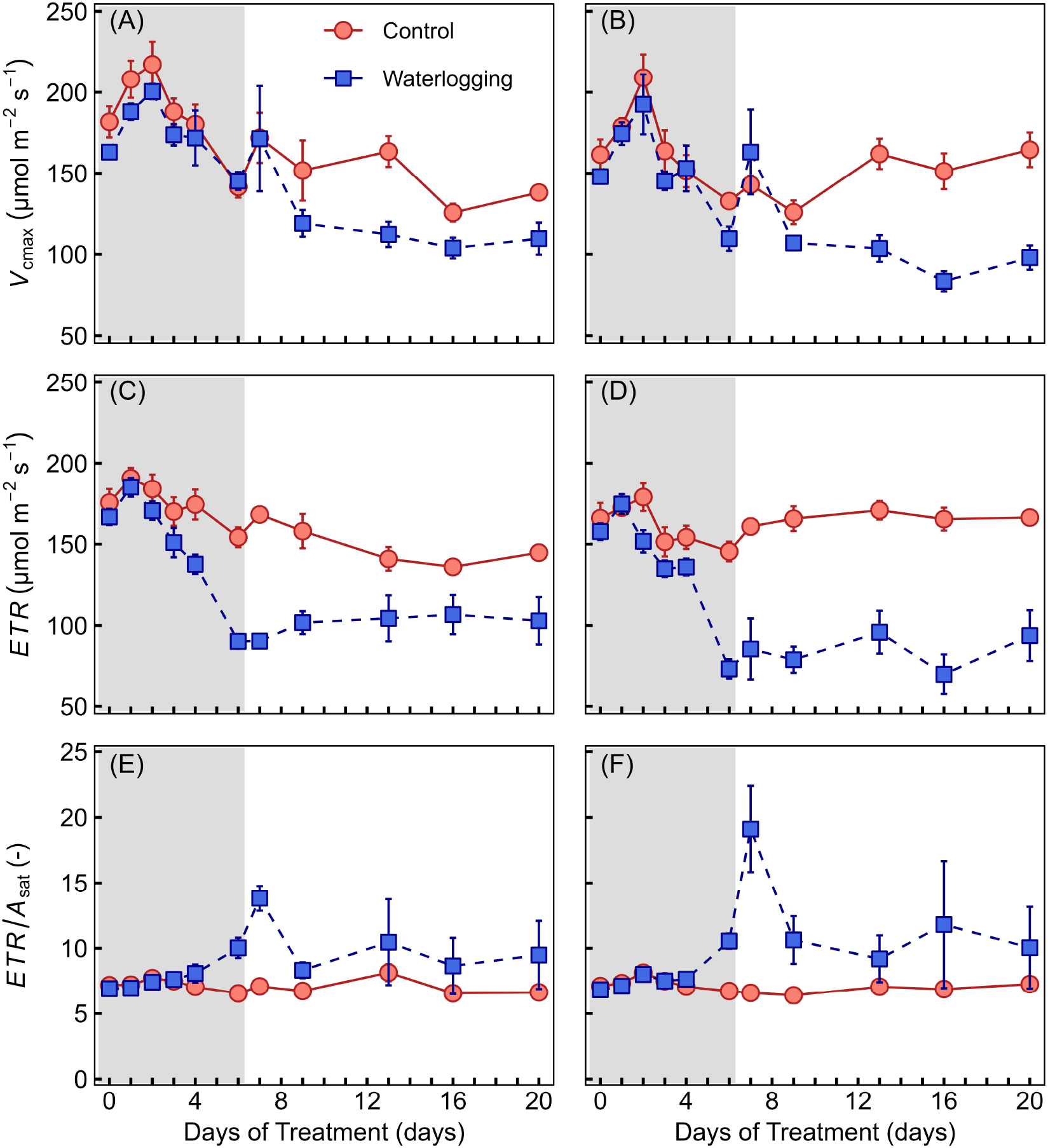
Temporal changes in maximum carboxylation rate based on CO_2_ concentration at the carboxylation site *V*_cmax_ (A, B), electron transport rate *ETR* (C, D), and the ratio of light-saturated photosynthetic rate and electron transport rate *ETR/A*_sat_ (E, F) during waterlogging and reoxygenation period. The red circles and solid line indicate the measured values in control, and the blue squares and dushed line indicate the measured values in waterlogging treatment. The respective rows indicate the soybean cultivar (i.e. A, C, E: Fukuyutaka, B, D, F: Iyo-daizu). Gray shaded area indicates the waterlogging period. Plots and error bars denoted means ± standard error (*n* = 4 to 5).

### 3.2 Relationship between photosynthetic rate and photosynthetic parameters

*A*_sat_ and *g*_sc_ and *g*_m_ showed a significant positive correlation during both soil waterlogging and reoxygenation periods, while *A*_sat_ during reoxygenation was smaller compared to that during waterlogging even under same *g*_sc_ or *g*_m_ in both cultivars (Fig. 3 (A) and (B)). *A*_sat_ and *C*_c_ showed a significant positive correlation during both soil waterlogging and reoxygenation period (Fig. 3 (C)). Similar to diffusional conductance, *A*_sat_ during reoxygenation was smaller compared to that during waterlogging even under same *C*_c_ in both cultivars. These results imply a limitation of *A*_sat_ by *V*_cmax_ during reoxygenation. Indeed, *V*_cmax_ during reoxygenation was relatively smaller than that during waterlogging, and a significant positive correlation between *A*_sat_ and *V*_cmax_ was found only under waterlogging (Fig. 3 (D)).

**Fig. 3:**
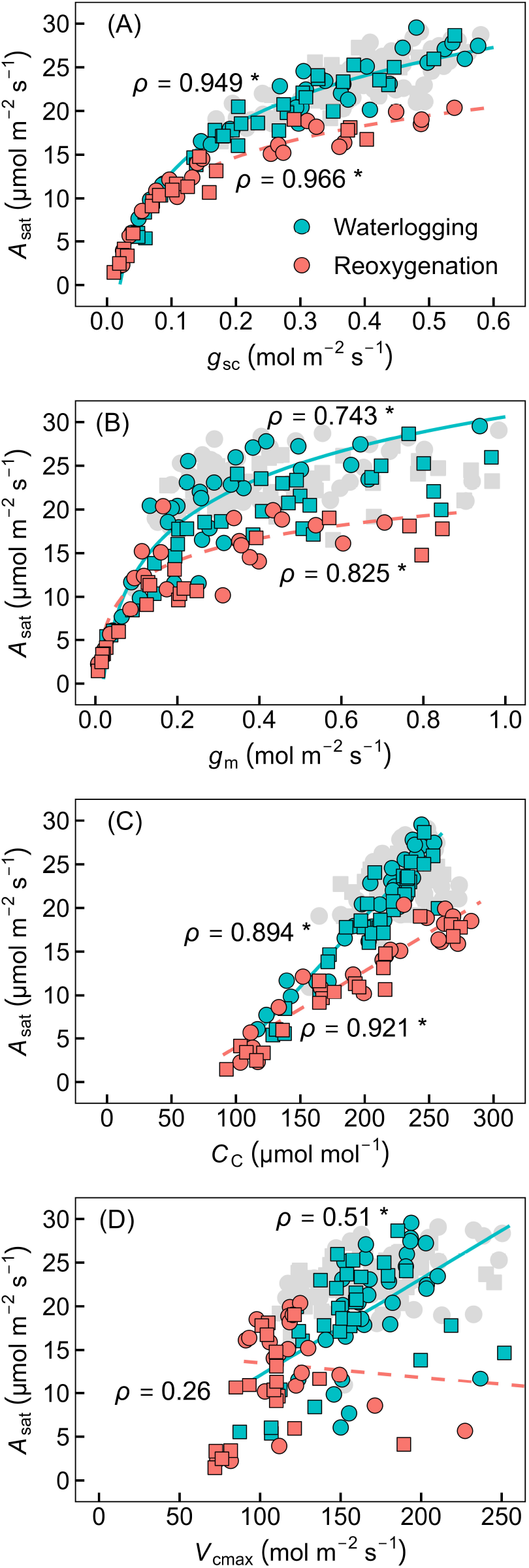
Relationship between light-saturated photosynthetic rate *A*_sat_ and stomatal conductance to CO_2_ *g*_sc_ (A), mesophyll conductance *g*_m_ (B), CO_2_ concentration at the carboxylation site *C*_c_ (C), and maximum carboxylation rate based on CO_2_ concentration at the carboxylation site *V*_cmax_ (D) in waterlogging treatment. Blue and red plots indicate the measured values during waterlogging and reoxygenation, respectively. Circle and square plots indicate the measured values of Fukuyutaka and Iyodaizu, respectively. Grey plots indicate the measured values in control. The curves indicate respective regression lines. The *Spearmann*’s rank correlation coefficients (ρ) and statistical significance are also shown (*: *p*<0.001).

### 3.3 Photosynthetic limitation during waterlogging and reoxygenation

In both cultivars, the total limitation (i.e., *S*_L_ + *M*_L_ + *B*_L_) of photosynthetic rate for the first 2 days after the onset of waterlogging was almost zero (Fig. 4). During the subsequent late-waterlogging period, the total limitation was sharply increased to 63.4% and 67.0% in Fukuyutaka and Iyodaizu, respectively, largely due to diffusional limitations (i.e., *S*_L_ + *M*_L_). In Fukuyutaka, although the total limitation decreased to 23.3% by the 6 days after reoxygenation (13 days of total treatment), it did not decrease any further until the end of the experiment (Fig. 4 (A)). This decrease in photosynthetic limitation was associated with a decrease in diffusional limitation. *B*_L_ increased to 20% by 6 days after reoxygenation and became the primary limiting factor for photosynthesis during the reoxygenation period. In Iyodaizu, the total limitation slightly decreased to 48.7% by the 6 days after reoxygenation (Fig. 4 (B)). This was due to an increase in *B*_L_ to 32.1%, as well as the maintenance of diffusion limitation at 16.6%, since stomatal conductance did not fully recover even during the reoxygenation period (Fig. 1 (D)).

**Fig. 4:**
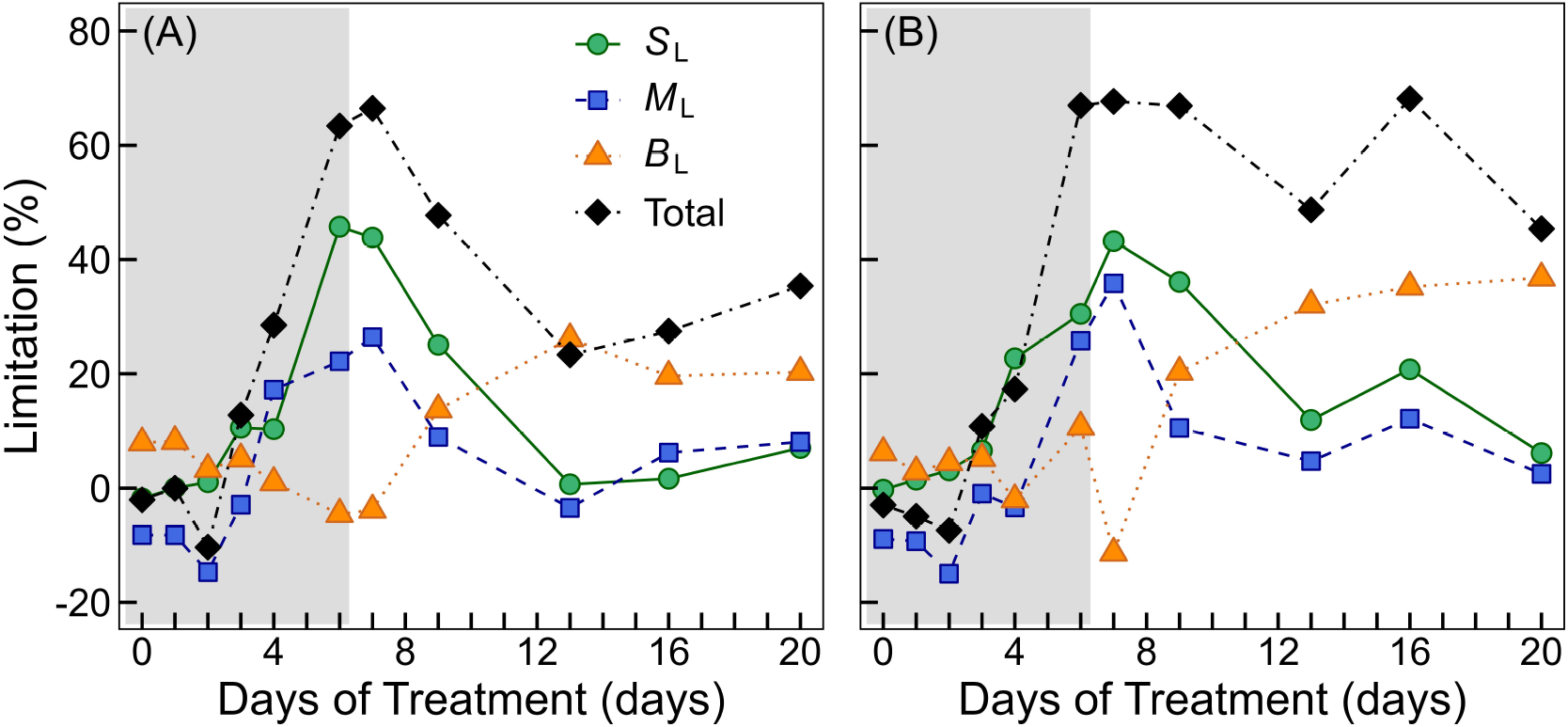
Temporal changes in limitations of light-saturated photosynthetic rate *A*_sat_ (%) for Fukuyutaka (A) and Iyo-daizu (B) during soil waterlogging and reoxygenation period in comparison with control. *S*_L_, *M*_L_, *B*_L_, and Total indicate stomatal conductance limitation (green circles), mesophyll conductance limitation (blue squares), biochemical limitation (orange triangle), and the sum of each limitation (black rhombus), respectively. Gray shaded area indicates the waterlogging period.

## 4. DISCUSSION

The present study provides data on leaf gas exchange and chlorophyll fluorescence under waterlogging and subsequent reoxygenation in two soybean cultivars and clarifies the extent to which photosynthesis is limited by stomatal conductance, mesophyll conductance, and the maximum carboxylation rate. To the best of our knowledge, this is the first attempt to apply quantitative limitation analysis (Grassi and Magnani, 2005) to waterlogging conditions. In both cultivars, *A*_sat_ started to decline significantly after 4 days of waterlogging and did not fully recover by two weeks of reoxygenation. Given that waterlogging lasting more than 5 days is frequently observed in both natural ecosystems and agricultural fields in humid regions (Kubota et al., 2023; Mercau et al., 2016; Villalobos-Vega et al., 2014), waterlogging stress can have a serious impact on plant growth. The primary limitation of *A*_sat_ was caused by leaf diffusional factors (a combination of stomatal and mesophyll conductance) during waterlogging and by biochemical factors during post-waterlogging reoxygenation. These findings highlight the need for comprehensive, time-course investigations of CO_2_ diffusion from the atmosphere to the carboxylation site, light-dependent reactions, and carbon assimilation reactions in the Calvin cycle to understand photosynthetic responses to waterlogging stress.

### 4.1 Photosynthetic limitation during waterlogging

In the context of waterlogging-induced photosynthetic limitation, the leaf stomatal factor is widely recognized (Garcia et al., 2020; Herzog et al., 2016; Pompeiano et al., 2019; Toral-Juarez et al., 2021). Here, we observed a significant decline in *g*_s_ during waterlogging in both cultivars and a subsequent limitation of photosynthetic rate by the stomatal factor. Leaf wilting was visibly observed during waterlogging, and thus the decline in *g*_s_ in our study likely reflects leaf water depletion, which can be attributed to a decrease in water uptake capacity and high evaporative demand (up to 3.4 kPa vapor pressure deficit in the greenhouse; Fig. S2). Many previous studies have demonstrated that waterlogging stress induces significant decrease in hydraulic conductance of both roots (Araki 2006; Else et al., 2001; Rodríguez-Gamir et al., 2011) and shoots (Haverroth et al., 2025), contributing to reduced leaf water potential in a broad range of plant species, including soybean plants (Garcia et al., 2020; Kubota et al., 2023; Kubota et al., 2025). However, some soybean cultivars maintain leaf water potential under waterlogging (Araki, 2006; Garcia et al., 2020) but still exhibited a reduced *g*_s_, indicating that leaf water status alone cannot explain stomatal closure. Although several chemical signals, such as abscisic acid, ethylene, and reactive oxygen species (ROS), have been proposed as additional triggers for stomatal closure under waterlogging (Else et al. 2009; Rodríguez-Gamir et al. 2011), the drivers of stomatal control remain unclear. Further studies that simultaneously monitor *g*_s_, leaf water status, and chemical signals are needed to elucidate the underlying mechanisms regulating stomatal aperture.

An increase in *C*_i_ in response to a decrease in *A* and *g*_s_ was observed in some plant species, such as *Arundo donax* L. (Pompeiano et al., 2019), *Picea sitchensis* (Black et al., 2005), *Brassica napus* and *Pisum sativum* (Ploschuk et al., 2018), suggesting a non-stomatal photosynthetic limitation (Farquhar and Sharkey, 1982). In soybeans, *C*_c_ decreased along with *A*_sat_, *g*_sc_, and *g*_m_ after 4 days of waterlogging (Fig. 1), and thus *g*_m_ also limited the *A*_sat_ by 22.2% and 25.8% during waterlogging (Fig. 4). While many studies have suggested a significant impact of *g*_m_ on photosynthetic limitation under soil drought (Egea et al., 2011; Flexas et al. 2009; Galle et al., 2009), studies on photosynthetic limitations under soil waterlogging conditions have mainly focused on stomatal and biochemical factors (Else et al., 2009; Garcia et al., 2020; Toral-Juarez et al., 2021), with only a few studies investigating *g*_m_ responses (Ploschuk et al., 2018; Pompeiano et al., 2019). Therefore, the mechanistic understanding of *g*_m_ control under waterlogging remains limited. *g*_m_ responds to multiple environmental and physiological factors, including CO_2_ concentration, leaf temperature, and leaf water status (Evans, 2021; Flexas et al., 2008). In the present study, measurements were taken under constant light intensity and CO_2_ concentration, and *g*_m_ was standardized to 25°C leaf temperature using a modified Arrhenius equation. Therefore, the decline in *g*_m_ under waterlogging would be mainly related to leaf water stress. These findings highlight the importance of evaluating *g*_m_ as well as *g*_s_ to accurately assess the limiting factors of the photosynthetic rate under waterlogging, as ignoring *g*_m_ can lead to an underestimation of the diffusional limitations (Flexas et al. 2009; Galle et al., 2009; Grassi and Magnani, 2005).

### 4.2 Photosynthetic limitation during reoxygenation

Substantial legacy effects of waterlogging on root metabolism (da-Silva et al., 2020), plant hydraulics (Haverroth et al., 2025), leaf gas exchange (Toral-Juarez et al., 2021), and plant growth (Ploschuk et al., 2018) have been observed across a broad range of plant species during post-waterlogging reoxygenation. Many previous studies have demonstrated that sudden exposure of roots to oxygen after prolonged waterlogging induces redox imbalance in roots and leaves (da-Silva et al., 2020; Garcia et al., 2020; Martins et al., 2024), which may inhibit the rapid recovery of leaf gas exchange after drainage (Toral-Juarez et al., 2021). The physiological responses to reoxygenation vary with plant species (Ploschuk et al., 2018), cultivars (Garcia et al., 2020; Toral-Juarez et al., 2021) and waterlogging stress intensity (Kubota et al., 2025; Pompeiano et al., 2019). In the present study, *A*_sat_ in both soybean cultivars could not recover completely through the approximately two-week reoxygenation period (Fig. 1). In addition, ETR/*A*_sat_ values steeply increased and exceeded 10.5 immediately after excess water drainage (Fig. 2 (E) and (F)). In non-stressed C_3_ plants, ETR/*A* values usually range from 7.5 to 10.5 (Flexas et al. 2002; Perera-Castro and Flexas, 2023), supporting the interpretation that soybeans would still be in a stressed state during the early stages of reoxygenation. Reoxygenation often induces oxidative stress, characterized by the accumulation of ROS in leaves (da-Silva and do Amarante, 2020; Garcia et al., 2020), the impairment of the electron transport chain (Posso et al., 2020), a decrease in leaf chlorophyll content (Da-Silva and do Amarante, 2020; Martins et al., 2024), and finally a marked limitation of photosynthetic rate (Toral-Juarez et al., 2021).

During this period, *A*_sat_, *g*_sc_ and *g*_m_ in both cultivars showed little recovery (Fig. 1), resulting in leaf diffusional conductance still being the predominant limiting factor for the photosynthetic rate during the early stages of reoxygenation (Fig. 4). As declined plant hydraulic conductance during waterlogging cannot recover immediately during the reoxygenation process, continuous leaf water stress after drainage has been observed in some plant species (e.g. Haverroth et al., 2025; Toral-Juarez et al. 2021) including soybeans (Kubota et al., 2025). In addition, the accumulation of apoplastic ROS induces stomatal closure via activation of Ca^2+^-permeable channels in guard cells (Kollist et al., 2014; Qi et al., 2018). Therefore, diffusional limitations for photosynthesis during early reoxygenation may result from the combined effects of water stress and oxidative stress.

Notably, despite recovering *g*_sc_ and *g*_m_ after 7 days of reoxygenation, the photosynthetic rate in the waterlogging treatment was still limited (Fig. 3 (A) and (B)), which was attributed to a significant decrease in *V*_cmax_. Thus, photosynthetic limitation due to biochemical factors became dominant after 7 days of reoxygenation (Fig. 4). A similar decrease in *V*_cmax_ during reoxygenation was observed in some crop species, such as *Vigna unguiculata* L. (Olorunwa et al., 2022b), *Coffea. canephora* (Toral-Juarez et al., 2021), and *Triticum aestivum* (Yang et al., 2024). Previous studies have attributed the decrease in *V*_cmax_ to the expression of rubisco and rubisco activase genes (Kuai et al., 2014; Ma and Guo, 2014), which may be associated with leaf N deficiency (Herzog et al., 2016) and chlorophyll degradation (Ma and Guo, 2014; Olorunwa et al., 2022a). Overall, diffusional factors and biochemical reactions (maximum carboxylation rate) were the primary limiting factors for *A*_sat_ in soybean plants during early- and late-reoxygenation, respectively. Our findings highlight the importance of comprehensive measurements of limiting factors such as stomatal, mesophyll, and biochemical reactions, as photosynthesis is limited by multiple factors that change over time during reoxygenation. However, it should be noted that methodological bias in quantitative limitation analysis potentially leads to an underestimation of the biochemical limitation. Quantitative limitation analysis of photosynthetic rate is derived from the biochemical model of C_3_ photosynthesis (Farquhar et al. 1980), which is defined as the photosynthetic rate limited by the carboxylation of RuBP (ribulose-1,5-bisphosphate) or by RuBP regeneration. In the present study, CO_2_ concentration in the chamber was set to 350 μmol mol^−1^ so that RuBP carboxylation limited the photosynthetic rate. Although *C*_c_ was smaller than *C*_ctr_ in most measured data, RuBP regeneration still limited photosynthesis in some data. Considering that the current atmospheric CO_2_ concentration is approximately 420 μmol mol^−1^ and will increase in the future, further studies is important to evaluate the biochemical limitations caused by RuBP regeneration rate, *i*.*e*. maximum electron transport rate.

### 4.3 Different photosynthetic response to reoxygenation between cultivars

Sakazono et al. (2013) and Suematsu et al. (2017) reported that Iyodaizu is a waterlogging-tolerant cultivar because it maintains a high growth rate even under 7 days of waterlogging, whereas Fukuyutaka exhibits strong growth inhibition. Interestingly, in the present study, both cultivars showed similar photosynthetic responses, with a significant decrease in *A*_sat_ during waterlogging (Fig. 1 (A) and (B)). Together, these findings suggest that Fukuyutaka may prioritize maintaining nonstructural carbohydrate (NSC) reserves by limiting growth during stress, while Iyodaizu may sustain growth by consuming its NSC reserves. These trade-off relationships between growth and carbon storage under waterlogging were highlighted by Striker (2012) and confirmed in some previous studies (e.g. Camisón et al., 2020; Yang et al., 2019). The NSC reserves act as buffers that allow recovery after stress (Wang and Wang, 2025), including waterlogging (Kuai et al., 2014; Qin et al., 2013). Indeed, Fukuyutaka partially recovered its photosynthetic rate during reoxygenation, whereas Iyodaizu showed little recovery (Fig. 1 (A) and (B)). To elucidate cultivar-specific tolerance mechanisms, future research should integrate NSC dynamics, photosynthetic traits responses, and growth performance during stress and recovery.

## 5. CONCLUSION

We investigated the dynamic processes constraining photosynthesis during soil waterlogging and subsequent reoxygenation in two soybean cultivars, with contrasting growth strategies. In both cultivars, the photosynthetic rate significantly decreased after more than 4 days of waterlogging and did not recover completely by two weeks of reoxygenation. Diffusional limitations (combination of *g*_s_ and *g*_m_) dominated during waterlogging and persisted into the early phase of reoxygenation. Thereafter, biochemical limitations, caused by reduced *V*_cmax_, became the primary constraint during late reoxygenation. This result indicates that impaired Rubisco activity and associated biochemical processes may delay photosynthetic recovery. Therefore, the most effective targets for increasing carbon gain under waterlogging conditions are likely to be maintaining high *g*_s_ and *g*_m_ during waterlogging and high *V*_cmax_ during reoxygenation. Our findings demonstrate that photosynthesis is dynamically limited by multiple factors during waterlogging and reoxygenation, highlighting the importance of comprehensive measurements of gas exchange and carbon assimilation reactions.

## Abbreviations

*A*_sat_: net CO_2_ assimilation rate under saturated light condiitons
*C*_i_: substomatal CO_2_ concentration
*C*_c_: chloroplast CO_2_ concentration
*E*_a_: activation energy
ETR: photosynthetic electron transport rate
*g*_sc_: stomatal conductance for CO_2_
*g*_sw_: stomatal conductance for water vapor
*g*_m_: mesophyll conductance for CO_2_
*H*_d_: deactivation energy
*K*_m_: Michaelis–Menten constants
PPFD: photosynthetic photon flux density
*R*_d_: leaf respiration in the dark
ROS: reactive oxygen species
*S*_*L*_: stomatal limitation
*M*_*L*_: mesophyll conductance limitation
*B*_*L*_: biochemical limitation
*T*_leaf_: leaf temperature
*V*_cmax_: maximum carboxylation rate on the basis of chloroplastic CO_2_ concentration
α: leaf absorptance
β: electron partitioning between PSI and PSII
*Γ*\*: CO_2_ compensation point under non-photorespiratory conditions
Δ*S*: entropy factor
Φ□_PSII_: photochemical yield of PSC□

## Supplementary Materials

Table S1. Parameters used for calculation of mesophyll conductance and maximum carboxylation rate

Table S2. The minimum leaf conductance of two soybean cultivars for each treatment.

Fig. S1. Photographs of soybean plants used for the experiment.

Fig. S2. Time courses of micrometeorological conditions during experimental period.

Fig. S3. The relationship between leaf temperature and mesophyll conductance and maximum carboxylation rate.

Fig. S4. The relationship between estimated *V*_cmax_ by *A*-*C*_i_ curve method and that by one-point method.

## Acknowledgement

We are grateful to Professor Toyoaki Anai for introducing the waterlogging-tolerant soybean cultivar (Iyodaizu) and providing its seeds used in present experiment. This research was financially supported by Grants-in-Aid for Scientific Research (JSPS KAKENHI grant number 23K19318) from the Japan Society for the Promotion of Science and by the University Reform and Revitalization System in Kyushu University.

## Author contributions

SK: conceptualization; SK, GY: methodology; SK, GY, TS: investigation; SK, GY: Formal analysis; SK, GY, TS, DY: Resources; SK: writing - original draft; GY, TS, DY, TH: writing - review & editing; SK, TH: funding acquisition

## Data availability

Data will be made available on request.

